# Integrative Gene and Isoform Co-expression Networks Reveal Regulatory Rewiring in Stress-related Psychiatric Disorders

**DOI:** 10.1101/2025.03.24.640781

**Authors:** Ghalia Rehawi, Jonas Hagenberg, BeCOME study group, OPTIMA study group, Philipp G. Sämann, Lambert Moyon, Elisabeth Binder, Markus List, Annalisa Marsico, Janine Knauer-Arloth

## Abstract

Isoform-specific expression patterns have been implicated in stress-related psychiatric disorders like major depressive disorder (MDD), yet the extent of their involvement and their interrelationships remain unclear.

We constructed co-expression networks for individuals affected (n=210, 81% with depressive symptoms) and unaffected (n=95) by stress-related psychiatric disorders. We incorporated total gene expression (TE) and isoform ratio (IR) data and validated the inferred networks using advanced graph generation techniques.

Our analysis revealed distinct network topology and structure between the two groups. Investigation of the 127 shared hubs (degree >= 10) found that these hubs exhibit co-regulatory patterns unique to each network. The affected individuals’ network also contained 61 hub nodes with a minimum absolute fold increase of two in connectivity compared to the unaffected individuals’ network. Notably, 49% of these hubs showed evidence for association with psychiatric disorders. Gene Ontology enrichment analysis revealed distinct biological processes associated with hubs, such as mRNA processing for affected and immune response and cell adhesion for unaffected individuals. Enrichment analysis of GWAS loci further supported network-specific findings. Analysis of the isoform-specific nodes showed distinct protein-protein interactions compared to gene-level analysis.

This is the first study to demonstrate network-level differences in gene and isoform co-expression patterns between individuals with and without stress-related psychiatric disorders, with a particular focus on depressive symptoms. Our findings provide evidence for substantial rewiring of gene regulatory networks in affected individuals. Incorporating isoform-level data revealed a deeper level of complexity, highlighting the importance of considering isoform variations in understanding the molecular basis of these conditions.

## Introduction

Stress-related psychiatric disorders, such as major depressive disorder (MDD), anxiety disorders, and post-traumatic stress disorders (PTSD) share common pathophysiological, clinical, and biological characteristics and impose a significant burden on individuals and society ^1,2^. These conditions disrupt thinking, mood, and daily functioning, leading to diminished quality of life and often long-lasting disability. This burden extends to healthcare systems, where psychiatric disorders are a leading cause of disability and contribute to poor outcomes in physical diseases ^1^. MDD exemplifies the challenges in understanding and treating psychiatric disorders. As a highly polygenic disease, MDD is influenced by numerous genetic variants, and its high comorbidity with many other psychiatric disorders complicates its study ^3^. Cross-disorder psychiatric studies offer a valuable approach to investigating shared biological processes beyond phenotypic features ^4^.

Through Genome-Wide Association Studies (GWAS), the majority of disease-associated variants were found to be in non-coding regions, highlighting the importance of gene expression and splicing regulation in contributing to genetic risk. This has led to increased interest in studying the gene expression landscape and transcriptional regulation. Differential Expression Analysis (DEA) is an important tool that allows researchers to identify genes expressed at significantly different levels between two or more conditions. Using samples from brain and blood tissues, many studies have identified transcriptional dysregulation patterns in patients with psychiatric disorders ^5–10^, with multiple differentially expressed genes being shared across several psychiatric disorders ^7,8^.

To unravel the complex biology of psychiatric disorders, it is important to organize genes within their broader molecular system and pathways context. However, DEA often focuses on individual genes, potentially overlooking the complex interactions and regulatory relationships within biological systems. To this end, co-expression networks have emerged as a powerful tool. This approach involves constructing networks representing functional relationships between genes, where nodes represent genes and edges represent a correlation of expression patterns. Network methods allow researchers to identify key regulatory genes, and modules of functionally related genes, and link them to disease-related pathways, offering a more comprehensive view of the molecular mechanisms underlying psychiatric disorders ^11–13^.

Network approaches have been widely used to investigate the pathophysiology of psychiatric disorders ^12–16^. For example, studies have used network methods to explore gene interactions, identifying key hub genes and modules associated with MDD status from blood samples ^14,16^. However, most existing methods have focused on investigating gene-level interactions ^8,12,13,16^, disregarding the effect of post/co-transcriptional modification processes, including alternative splicing (AS). AS affects up to 95% of human genes ^17^, plays an important role in gene regulation, and contributes to the diversity and complexity of the proteome ^18–20^ by producing different isoforms of the same gene, with much research demonstrating that different isoforms of the same gene may have different or even opposing functions ^20–22^. Recently, considerable effort has been directed toward studying AS and splicing dysregulation in psychiatric disorders ^7,23–25^. For instance, an increase in the expression of specific isoforms of the neuregulin 1 receptor *ERBB4* has been reported in patients with schizophrenia ^26^. Another study identified differentially spliced genes, including splicing regulators, in patients with autism spectrum disorder (ASD) ^27^. Studies incorporating isoform-level data into differential expression and network analysis have revealed larger effect sizes and more informative disease-specific transcriptional profiles and biological signals often missed when focusing solely on gene-level expression ^7,28,29^. For example, in a cross-disorder study of ASD, SCZ, and BP, Gandal et al.^7^ demonstrated that isoform-level co-expression networks were more strongly associated with disease-specific GWAS loci than gene-level networks.

While these studies have highlighted the importance of isoform-level analysis in understanding psychiatric disorders, there remains a need for integrative approaches that combine both gene-level and isoform-level data in a single network framework. To address this need, we introduce an integrative network approach to compare and unravel the complex underlying biology between a network of affected individuals (AIN) with stress-related psychiatric disorders (n=210, 81% with depressive symptoms) and a network of unaffected individuals (UIN) (n=95). As in the work studying tissue-specific transcription and splicing by Saha et al. ^30^, we combine both total gene expression values (TE) and isoform ratios (IR) as two node modalities in our networks. Using advanced graph generation and embedding techniques, we validate that these networks capture biologically meaningful distinctions between the two groups. We compare the two networks to reveal differences in co-regulatory patterns both at gene and isoform levels. Additionally, we prioritize key genes and isoforms within the AIN that may play pivotal roles in disease pathways and serve as potential targets for therapeutics.

To elucidate the advantages of our network-based approach over current standard methods such as differential expression analysis, we perform DEA at both gene and transcript levels, followed by pathway and GWAS enrichment analyses on both DEA results and network findings. By constructing and comparing integrative gene and isoform networks for affected and unaffected individuals, we reveal changes in regulatory relationships and gain novel insights that are not captured from differential expression analysis alone.

## Materials and Methods

### 1. Samples Selection

This study included 336 Caucasian participants selected based on the availability of matching RNA sequencing (RNAseq) and phenotypic data from three cohorts recruited at the Max Planck Institute of Psychiatry in Munich: The Biological Classification of Mental Disorders (BeCOME) study (ClinicalTrials.gov: NCT03984084, ^31^), the Imaging Stress Test (IST) study, and The OPtimized Treatment Identification at the Max Planck Institute study (OPTIMA) (ClinicalTrials.gov: NCT03287362, ^32^). Individuals were assessed as affected/unaffected based on the Munich-Composite International Diagnostic Interview (DIA-X/M-CIDI) ^33,34^.

In total, our samples comprised 229 affected individuals (BeCOME: 122, OPTIMA: 107) who met either threshold or subthreshold DSM-IV-based DIA-X/M-CIDI criteria for any substance use, affective or anxiety disorder, including post-traumatic stress disorder and obsessive-compulsive disorder, within the last 12 months of enrollment. 186 of these participants had a (subthreshold) DSM-IV diagnosis of major depression or dysthymia. Unaffected individuals (BeCOME: 70, IST: 37) were defined as those without any DSM-IV-based DIA-X/M-CIDI diagnosis. However, to focus on a more specific set of psychiatric disorders, cases with pure nicotine dependence (without any other comorbid diagnosis) were excluded from the affected group and moved to the unaffected group, resulting in a total of 107 unaffected individuals.

All participants were assessed by the Beck Depression Inventory (BDI) II ^35^ and the Montgomery–Åsberg Depression Rating Scale (MADRS) ^36^. An overview of the sample characteristics is provided in Table 1.

**Table 1:**
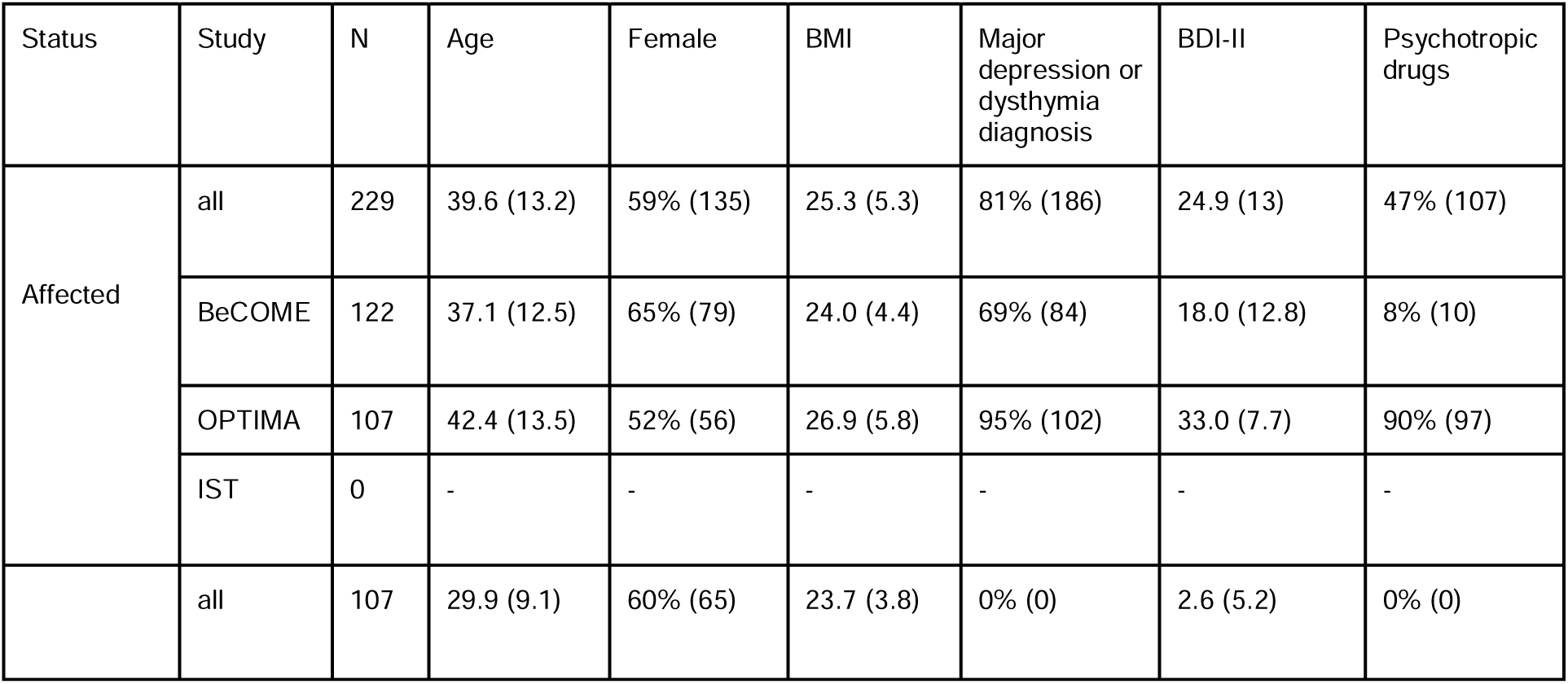
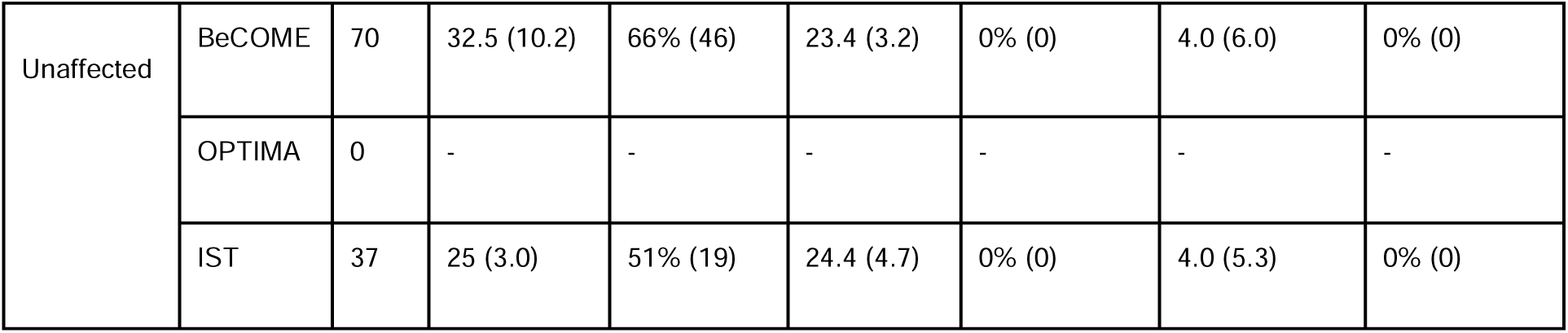
Cohort characteristics are presented as percentages with absolute numbers in parentheses for categorical variables, or as means with standard deviations in parentheses for continuous variables. Depression diagnoses, including both full and subthreshold diagnoses in the last 12 months, were determined using the Munich-Composite International Diagnostic Interview (M-CIDI). Depression severity was assessed using the Beck Depression Inventory-II (BDI-II). The column Psychotropic drugs refers to the percentage of subjects taking psychopharmacological medications during the study period. These medications include antidepressants, mood stabilizers, neuroleptics, tranquilizers, and herbal psychotropics.

The studies were approved by the ethics board of the Ludwig Maximilians University (approval BeCOME:#350–14, OPTIMA:#17–395, IST:#121-14) and conducted in accordance with the Declaration of Helsinki.

### 2. RNA extraction and sequencing

Blood samples were collected in the morning under fasted conditions. RNA was extracted from peripheral blood mononuclear cells (PBMCs) from OPTIMA and BeCOME cohorts and stored at the MPI biobank. Ribosomal RNA (rRNA) was depleted to enrich for messenger RNA (mRNA) and improve the detection of other transcripts using RiboCop rRNA Depletion Kits. Libraries were prepared with the Lexogen CORALL total RNA-Seq V1 Library Prep Kit and sequenced on a NovaSeq 6000 (Illumina, San Diego, USA) with a target depth of 30 million reads per sample, as previously described in more detail in ^37^. RNA was extracted from whole blood samples from the IST cohort. rRNA was depleted with RiboCop and libraries were prepared with the Lexogen CORALL total RNA-Seq V2 Library Prep Kit. Sequencing was performed on a NovaSeq 6000 in a separate batch with a target depth of 15.6 million reads per sample. Raw and processed sequencing data have been deposited in GEO under accession numbers GSE289144 and GSE289146.

### 3. RNA-seq alignment and QC

Paired-end FASTQ files were aligned against the GRCh38.p12 primary assembly using the GENCODE v31 annotation ^38^ with STAR aligner v2.7.7a. Alignment was performed using the option quantMode = TranscriptomeSAM, following protocol 7 of Dobin and Gingeras ^39^, which involves generating a transcriptome index and using it for alignment and quantification to produce output in transcriptomic coordinates. Gene and transcript-level expression were then quantified using RSEM v1.3.3 ^40^ for paired-end reads.

Gene and transcript-level reads were filtered for unwanted sequences using Cutadapt ^41^ v2.10. Zero-length reads were removed and only those with a count ≥ 10 in at least 95% of samples were retained, resulting in 9777 genes and 11427 transcripts.

Cell type deconvolution was calculated using Granulator v1.2.0 ^42^ and the LM22 reference matrix ^43^. Principal components (PCs) of the cell type proportions were calculated for inclusion in downstream models (see Supplementary Methods).

To account for confounding effects from different sequencing runs, we first corrected the gene and transcript-level data for the sequencing run using the ‘removeBatchEffect’ function from the limma R package ^44,45^. Subsequently, we performed Surrogate Variable Analysis (SVA) ^46^ to identify additional hidden batch effects. The first SV (sv1) was strongly correlated with the first principal component of gene expression (pc1), GC content, and the total number of filtered read pairs (total_pairs). Since these technical factors could confound our analysis, we further corrected gene and transcript-level data for GC content and total_pairs. We also removed the first five principal components of cell type proportions due to the different cell type composition across our samples (Supplementary Methods, Fig.S1), as well as the effect of sex, age, body mass index (BMI), as these biological factors could also introduce unwanted variation. Finally, we removed genes and transcripts with negative values due to the subtraction of the modeled biological and technical effects, resulting in 7394 genes and 7334 transcripts, which we used for differential gene and transcript analysis.

We removed technical and biological covariates to ensure that network inference primarily reflects the intrinsic differences between the affected and unaffected groups, preventing the network structure from being influenced by unrelated variations. For consistency, we used the same batch-corrected data for both differential expression analysis and network analysis.

### 4. Differential gene expression and transcript expression analysis

We carried out differential gene expression (DGE) and differential transcript expression (DTE) analysis on the corrected data using the limma-trend method ^44,47^. For this purpose, we transformed the corrected gene and transcript count data to log2-counts per million values (logCPM) and fitted the linear models using the functions lmFit and eBayes from limma. The design matrix included diagnosis (affected/unaffected) as the main factor of interest. Significant genes and transcripts were identified at an FDR of 5%.

### 5. Isoform ratios

Changes in the expression values of isoforms are influenced by various splicing regulatory processes and were shown to be associated with many diseases ^26,48–50^. Depending on the study objective, isoform-level information can be represented and modeled in several ways, including absolute isoform counts, isoform proportions, or isoform ratios relative to their parent genes ^19,28,30^. Similar to the work of Saha et al. ^30^, we modeled isoform abundances as ratios. Isoform ratios normalize transcript expression to the overall gene expression, reducing biases from gene-level variability and highlighting isoform-specific changes. The isoform ratio (IR) for each transcript was computed by dividing the corrected and logCPM-transformed transcript counts by the corresponding gene counts. Mapping isoforms to their genes was done using GENCODE V31 annotation ^38^. We removed transcripts whose genes had been filtered at previous steps. When none of the isoforms of a gene were expressed (0/0 divisions), the mean of the IR across all samples was taken ^30^. Hence, the final dataset for network inference includes 7394 genes (TEs) and 7097 isoform ratios (IRs).

### 6. Regulatory network inference for affected and unaffected individuals

Before constructing the networks for the affected and unaffected individuals, we performed principal component analysis (PCA) ^51,52^ on the logCPM gene count and the IR data separately. Outliers were identified based on a visual inspection of the first two principal components from the PCA, with a focus on data points that deviated by more than 2 standard deviations from the mean on either PC1 or PC2. A total of 31 outliers (19 affected and 12 unaffected) were removed. This resulted in 210 affected and 95 unaffected individuals being included in the network inference.

For both the affected individuals’ and unaffected individuals’ networks, we used ARACNE (Algorithm for the Reconstruction of Accurate Cellular Networks) ^53^, an information theoretic-based method designed for the reverse engineering of regulatory networks. ARACNE uses mutual information (MI) to identify potential interactions and the data processing inequality (DPI) to remove indirect relationships ^54^ (see Supplementary Methods). We used both gene expression and isoform ratios as input for ARACNE and removed edges connecting features of the same gene to reduce bias ^30^.

The initial results showed that the UIN has twice as many edges compared to the AIN with lower MI values of the inferred edges. This is likely due to a reduced inference power caused by the smaller sample size in the unaffected group (see Supplementary Results and Fig.S3). To enable meaningful and fair comparison, we applied a threshold based on the median MI value of each edge type in the affected network (Fig.S4), resulting in two networks with similar statistics (Table 2). Details on the ARACNE algorithm and thresholding procedure are provided in Supplementary Methods and Results.

**Table 2:** Overview of the number of nodes and edges as well as the number of edges for each edge type in the affected individuals and unaffected individuals’ networks. Values are shown before (top row) and after (bottom row) the removal of edges with low mutual information (MI) values.

| Network | #Nodes | #Edges | #TE-TE edges | #TE-IR edges | #IR-IR edges |
| --- | --- | --- | --- | --- | --- |
| Affected-Individuals (AIN) | 14,300 | 21,324 | 11951 | 1946 | 7427 |
|  | 7996 | 10702 | 5997 | 983 | 3722 |
| Unaffected-Individuals (UIN) | 14,447 | 40,676 | 24199 | 6993 | 9484 |
|  | 8272 | 11120 | 4837 | 1382 | 4901 |

### 7. Validation of topological differences between the constructed networks

In this analysis, we assessed the robustness of the network inference step in capturing group-specific biological processes reflected in a distinct topological structure. To this end, we leveraged ARROW-Diff ^55^, a novel approach for efficient large-scale graph generation. ARROW-Diff incorporates two key components in an iterative procedure to generate graphs that closely resemble an input ground truth graph in terms of structural properties. The first component is an auto-regressive random walk-based diffusion model, which learns the generative process of random walks sampled from the input ground truth graph. This model captures the original network’s structural characteristics and local connectivity patterns. The second component is a Graph Convolutional Network (GCN) ^56^, which is trained to predict the validity of the proposed edges from the first component.

Utilizing ARROW-Diff, we generated 100 graphs for each network (affected and unaffected individuals’ networks). These simulated graphs capture the structure of the corresponding input graph in terms of various graph statistics, including global clustering coefficient, triangle count, assortativity, and other relevant graph metrics on which the graph generation is evaluated. By generating simulated networks that resemble each of the affected/unaffected individuals’ networks, we introduce variability/noise to the networks inferred by ARACNE. This is because ARROW-Diff can add or remove edges from the generated graph while maintaining the intrinsic graph structure.

We hypothesize that if the network inference captures group-specific properties reflected in distinct topological structures, the simulated graphs should also exhibit such distinctiveness between the two groups, even with the introduced variability. To analyze the structural similarities and differences between the generated graphs, we employed a two-step dimensionality reduction approach. First, we embedded the 200 generated graphs using Graph2Vec ^57^ with an embedding dimension of 128 in order to capture the structure of such large graphs. This step transforms each graph into a high-dimensional vector representation, capturing its topological features. Subsequently, we applied a PCA ^51^ to map these high-dimensional embeddings into a lower-dimensional space, facilitating visualization and analysis of the clustering patterns among the embedded graphs.

### 8. Functional enrichment

#### 8.1. Enrichment analysis of biological processes and psychiatric risk

We conducted an enrichment analysis of the DE genes, DE transcripts, and master hubs’ neighbors within the networks of affected and unaffected individuals using enrichGO from ClusterProfiler ^58^ v4.12.6. This analysis was based on Gene Ontology (GO) biological processes ^59,60^. To enhance the clarity and interpretability of the results, we applied the simplification process from ClusterProfiler with default parameters to remove redundancy among enriched GO terms, focusing on the most representative biological processes. Furthermore, using the Generalized Gene-Set Analysis of GWAS Data (MAGMA) ^61,62^, we assessed the enrichment in genes carrying single nucleotide polymorphisms (SNPs) with genome-wide association to the following traits: Attention Deficit Hyperactivity Disorder (ADHD), Autistic Spectrum Disorder (ASD), Bipolar Disorder (BP), Major Depressive Disorder (MDD), Post-traumatic Stress Disorder (PTSD), and a Psychiatric Cross Disorder phenotype. To provide a comparative baseline, we included a GWAS for Height as a control trait. To run MAGMA, we used the NCBI gene location file build 37 and the 1,000 Genomes reference data file for SNP locations.

#### 8.2. Gene-disease association analysis for psychiatric disorders

To investigate the associations between genes and psychiatric disorders, we used the DisGeNET resource, a comprehensive platform integrating information on gene-disease associations from various expert-curated databases, GWAS catalogs, animal models, and scientific literature ^63^. We queried DisGeNET for a wide range of mental and psychiatric conditions, including anxiety disorders, major depressive disorder, and various substance abuse and dependency disorders (Table S1). To ensure the reliability of our analysis, we applied a filtering criterion, considering only gene-disease associations (GDAs) with a score greater than 0.4. We used gene names of both genes and genes of corresponding isoforms for all enrichment analyses according to the GENCODE V31 annotation.

### 9. Network annotation

We compiled an extensive list from multiple resources to annotate the nodes within the networks for known transcription and splicing regulators. For splicing factors, we integrated data from SpliceAid-F, a curated database of human splicing factors and their RNA binding sites ^64^, which provided 67 splicing factors. Additionally, we incorporated a collection of 277 genes involved in pre-mRNA splicing from ^65^, and 406 splicing factor genes from ^66^. For transcription factors, we leveraged the TFLink resource ^67^, a gateway for transcription factor-target gene interactions. This integration resulted in a final compilation of 1,606 known transcription factors and 517 known splicing factors (Table S2). To maintain consistency and facilitate cross-referencing, we utilized gene names for the annotation process throughout our analysis.

## Results

### 1. Differential expression analysis reveals distinct gene and transcript-level dysregulation

After adjusting for biological variables (sex, age, BMI, and cell type composition), and technical variables (sequencing run, GC content, and total read pairs), we performed differential gene expression (DGE) and differential transcript expression (DTE) analyses incorporating both total gene expression counts (n=7394 genes) and transcript expression counts (n=7334 transcripts) from 229 affected and 107 unaffected individuals (see Methods). Our DE analyses identified 450 differentially expressed genes (36% up-regulated) and 269 differentially expressed transcripts (30% up-regulated) at an FDR of 5% (Fig.2c,d and Tables S3, and S4). Notably, our DTE analysis identified 104 transcripts that did not show DGE (Fig.2a,b and Fig.S2), consistent with previous findings ^7,28,68^.

**Figure 1:**
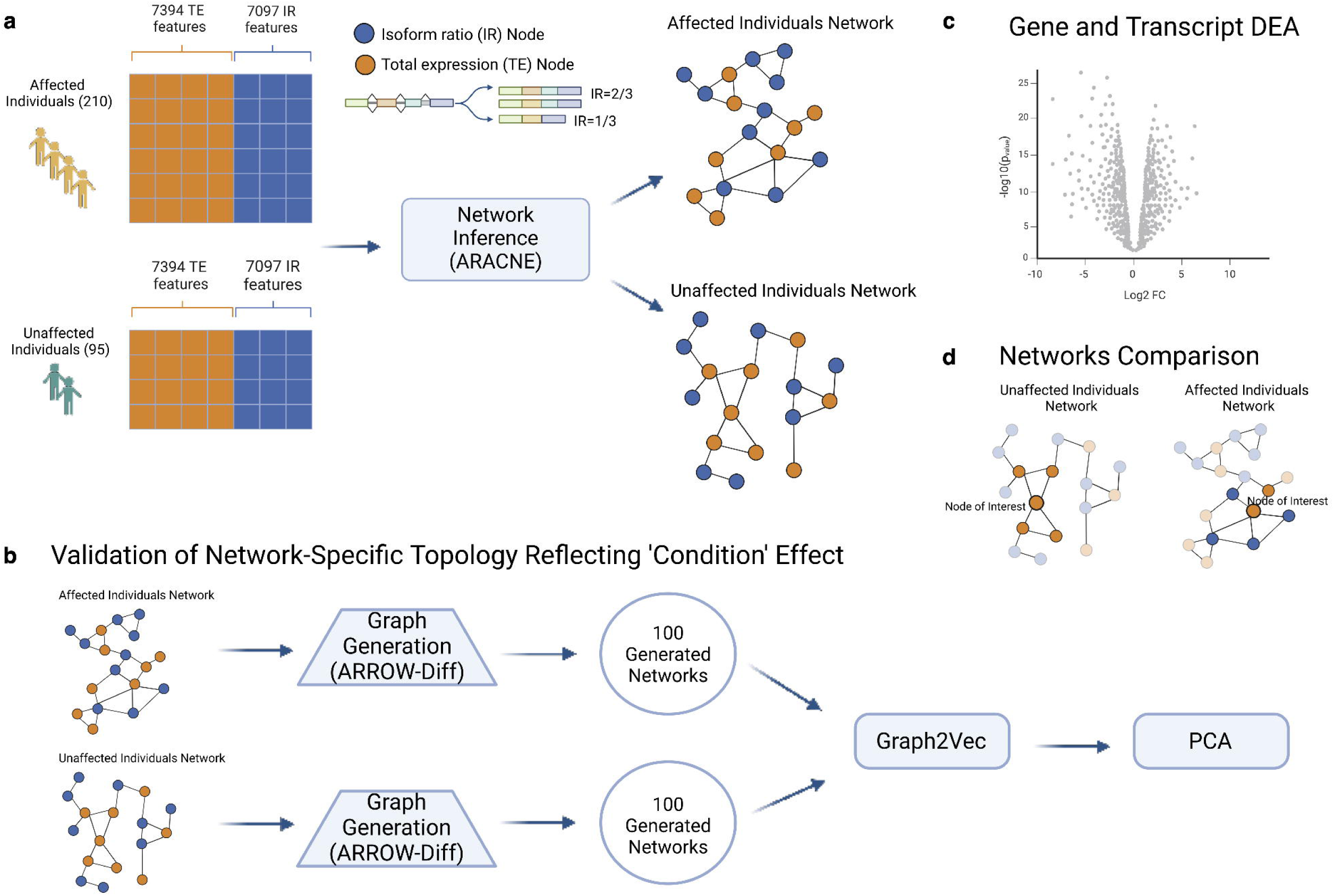
**a** Networks for affected (210) and unaffected (95) individuals were built using ARACNE network inference by incorporating both total gene expression values (TEs) and isoform ratios (IRs). **b** We employed graph AI techniques including the graph generation approach ARROW-Diff and graph embedding approach Graph2Vec to validate the distinct and network-specific topology for the inferred affected and unaffected individuals’ networks. **c** To elucidate the advantages of our network-based approach over traditional methods such as differential expression analysis, and to underscore the advantage of incorporating transcript level data, we performed DEA at both gene and transcript levels. **d** Affected and unaffected individuals’ networks were compared to elucidate the rewiring of co-regulatory relationships and differences in biological pathways and processes reflecting that these networks capture biologically meaningful distinctions between the two groups.

**Figure 2:**
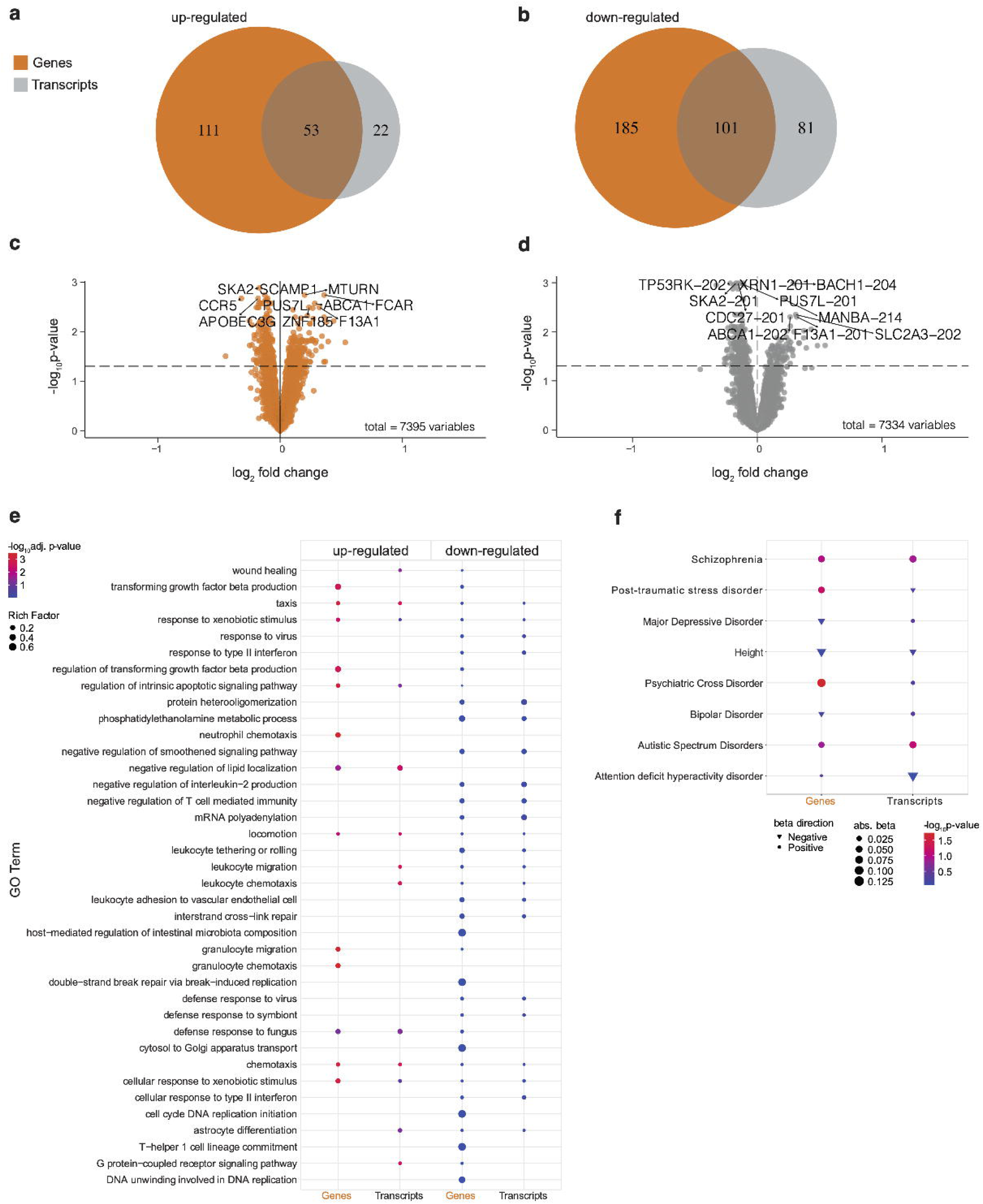
**a** Venn diagram of up-regulated genes and transcripts showing 53 up-regulated genes found both at gene and transcript level, and 22 genes corresponding to 23 up-regulated transcripts found only at the transcript level. **b** Venn diagram of down-regulated genes and transcripts showing 101 down-regulated genes found both at gene and transcript level, and 81 genes corresponding to 81 down-regulated transcripts found only at the transcript level. **c**,**d** Volcano plots of differential genes and transcripts at 5% FDR with the names of the top 5 (based on FDR) up and down-regulated genes and transcripts. **e** Enrichment in GO biological processes showing the top 10 (based on BH adj p-values) enriched processes for up and down-regulated genes and transcripts. Enrichment values, even if not significant, were filled for each process and for every set if available otherwise left empty (no dots). **f** Enrichment of DE genes and transcripts in genes that carry SNPs associated with the GWAS traits schizophrenia, post-traumatic stress disorder, major depressive disorder, bipolar disorder, autistic spectrum disorder, attention deficit hyperactivity disorder, and a psychiatric cross-disorder GWAS. The GWAS trait of Height was used as a baseline for comparison.

To identify pathways and biological functions relevant to the differentially expressed genes and transcripts, we carried out enrichment analysis of differentially expressed genes (DEGs) and differentially expressed transcripts (DETs) using ClusterProfiler ^58^, which revealed distinct biological processes associated with upregulated genes versus upregulated transcripts. Upregulated transcripts were uniquely enriched (Benjamini-Hochberg (BH) correction, p < 0.05) in leukocyte chemotaxis and leukocyte migration, both related to immune system processes. On the other hand, upregulated genes showed enrichment (BH, p < 0.05) in a different set of immune processes, including granulocyte chemotaxis, neutrophil chemotaxis, and granulocyte migration.

Neither downregulated genes nor downregulated transcripts exhibited significant enrichment in any GO biological processes. Figure 2e presents the top 10 enriched processes for DEGs and DETs, with a comprehensive table of enrichment results available in Table S5.

To further investigate the additional layer of information provided by transcript-level data in this context, we examined enrichment in genes carrying GWAS single nucleotide polymorphisms (SNPs) associated with psychiatric disorders using MAGMA ^61,62^. While differentially expressed genes captured more cross-disorder-related genes (β = 0.13, p = 0.038), differentially expressed transcripts showed a larger effect size for MDD-related SNPs (β = 0.01, p = 0.7) compared to differentially expressed genes (β = −0.03, p = 0.7). A similar pattern was observed for BP (see Fig.2f). The complete results of this analysis can be found in Table S6. Of all DEGs and DETs, we prioritized a list of genes that showed previous associations with psychiatric disorders. To this end, we intersected all differential genes and transcripts (553 unique genes) with a self-curated list of known psychiatric disorder-related genes (see Gene-disease association analysis in Methods section 8.2), yielding 53 genes, 15 of which are associated with depression (Table S7).

### 2. Construction and Validation of Co-expression Networks

To investigate differences in co-expression patterns of genes and isoforms between individuals affected (n=210, 81% with depressive symptoms) and unaffected (n=95) by stress-related psychiatric disorders, we constructed a co-expression network for each group and compared them. We employed the information theoretic-based network inference approach ARACNE ^53^ (see Methods) using both corrected total gene expression (TE) values (n=7394 genes) and isoform ratios (IR, n=7097 ratios). To select only high-confidence edges in both networks, we applied thresholding based on the MI values (see Methods). The resulting affected individuals’ network consists of 7996 nodes and 10,702 edges, while the network for unaffected individuals consists of 8272 nodes and 11,120 edges (Table 2), with a similar number of TE and IR nodes (Fig.3a).

**Figure 3:**
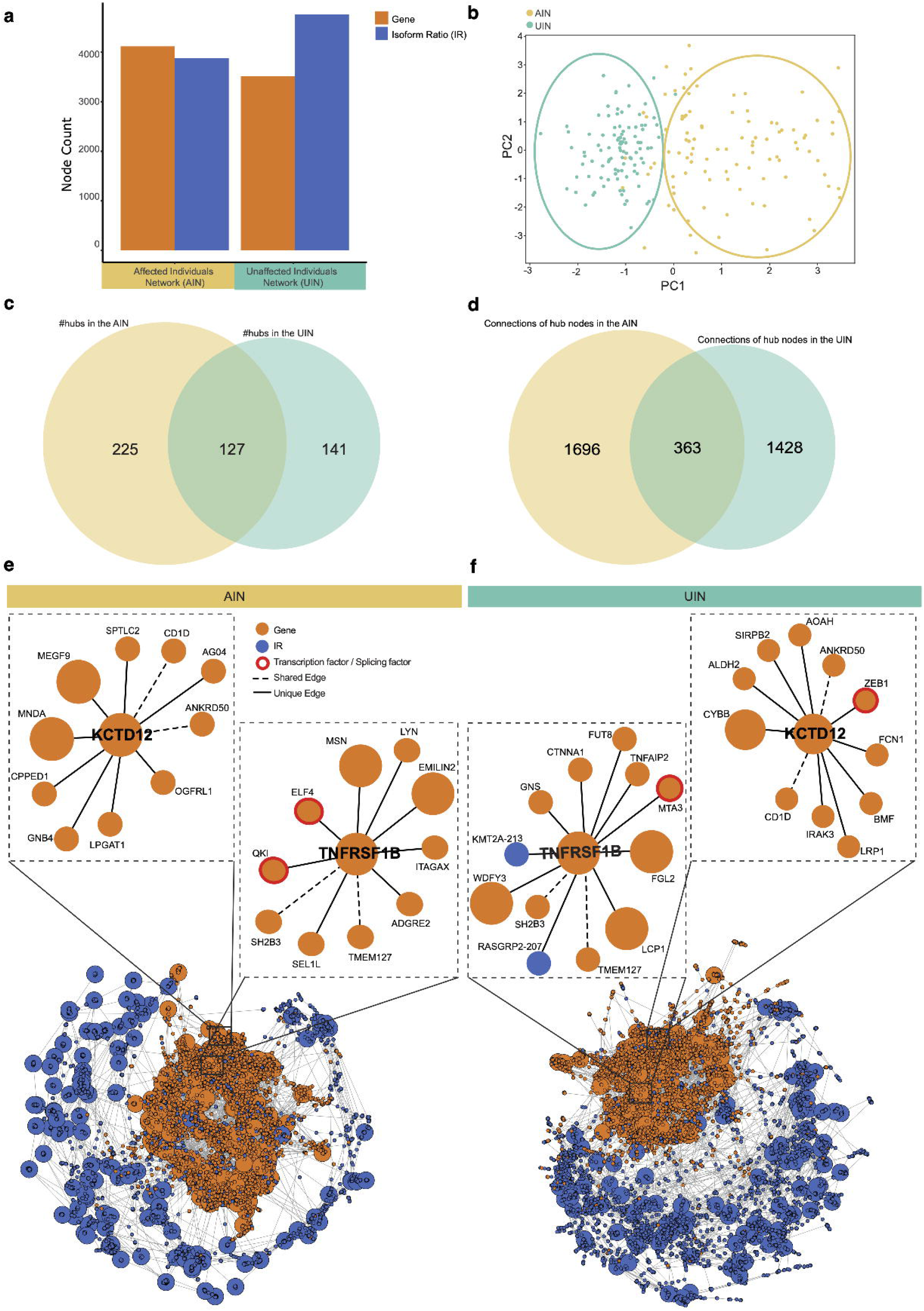
**a** Number of all TE and IR nodes in the two networks after filtering and thresholding. **b** PCA depicting a separate clustering of the Graph2vec embeddings of the 100 simulated UINs from the embeddings of the 100 simulated AINs validating the condition-specific network topology. **c** Venn diagram showing the number of hubs with degree >= 10 in both networks and their overlap. **d** Venn diagram showing the number of distinct connections of the 127 common hubs (degree >=10) in the two networks and their overlap. **e**,**f** First-order neighbors of two common hub genes associated with psychiatric disorders *KCD12* and *TNFRSF1B* are vastly different in the two networks, including distinct connections (solid lines) to regulatory transcription and splicing factors.

To ensure the robustness of the network inference in capturing biological differences between affected and unaffected individuals, we employed graph generation and embedding techniques (see Methods). Using ARROW-Diff ^55^, we created 100 simulated graphs for each network. We then investigated the structural similarities between the generated graphs by embedding them using Graph2Vec technique ^57^ and then mapping the embeddings into a lower-dimensional representation using PCA. Comparison of these simulated networks revealed a clear separation between the two groups’ networks (R^2^ = 0.99 linear regression score, Fig.3b). This quantitative assessment of the network topology distinctiveness between groups ensures robust inference and suggests that these networks reveal disease-specific patterns relevant to psychiatric diseases beyond noise.

### 3. Differential network analysis reveals distinct co-expression patterns associated with stress-related psychiatric diseases

To pinpoint key differences between the networks of affected and unaffected individuals, we focused on 1) investigating how common hub nodes differ in terms of connectivity patterns between the two networks, and 2) identifying key drivers of the underlying network-specific biology by investigating master hub nodes. To ensure that connectivity patterns are indeed distinct to each network, we only considered an edge to be distinct/unique to a network, if it is absent in the unthresholded version of the other network. By systematically comparing the connection patterns in the two networks, we aimed to uncover distinct patterns linked to psychiatric diseases both at the gene and isoform levels.

#### 3.1. Common hub nodes show distinct connection patterns between the affected and unaffected individuals’ networks

Hub nodes (degree ≥ 10), often central to biological processes and regulatory functions, were identified in both networks. The AIN contained 352 hubs (Table S8) and the UIN 268 (Table S9), with 127 hubs (degree ≥ 10) common to both networks, (Fig.3c, Table S10). These common hub nodes showed a distinct connection pattern in each network. Specifically, in the AIN, 1,696 of all common hubs’ connections (82.4%) were unique. Similarly, 1,428 (79.7%) of all common hubs’ connections in the UIN were unique. Only 363 edges were shared between the two networks (Fig.3d).

Of the 127 common hub nodes, 22 genes and genes of corresponding transcripts showed associations with psychiatric disorders based on the DisGeNet resource ^63^. Figure 3e,f depicts two of these genes, the Potassium Channel Tetramerization Domain Containing 12 (*KCTD12*), which is associated with bipolar disorder ^69^, and the TNF Receptor Superfamily Member 1B (*TNFRSF1B*) associated with depression ^70^, exhibiting substantially different connection patterns with only 2 edges common to both networks for the two genes. To elucidate the functional implications of these co-regulatory changes, we used the Domain Interaction Graph Guided ExploreR tool (DIGGER) ^71^ to investigate protein domain-level interactions. This analysis revealed that *TNFRSF1B*, which is co-regulated with TNF Alpha Induced Protein 2 (*TNFAIP2*) exclusively in the UIN (Fig.3f), interacts with the TNF alpha protein through the domain PF00020. Notably, TNF alpha has been implicated in MDD through its role in blood-brain barrier dysfunction ^72^. Furthermore, multiple studies have reported higher TNF levels in patients with depression ^73^. This suggests that a dysfunction in the protein domain PF00020 of TNF alpha, potentially leading to altered TNF signaling, may play a role in the pathology of psychiatric disorders. Overall, the divergence in connectivity patterns among common hubs suggests a potential rewiring of regulatory interactions in the context of stress-related psychiatric disorders.

#### 3.2. Master hub nodes reveal genes and transcripts relevant to stress-related psychiatric disorders in the affected individuals’ network

Our analysis revealed 61 master hub nodes in the AIN exhibiting substantial degree shifts, characterized by a minimum absolute fold increase of two in connectivity compared to the UIN, and a degree of at least 10. Of these, over half (n = 36) were isoform ratio nodes, with six also showing differential expression, including Complement C5a Receptor 1 (*C5AR1*), Caveolae Associated Protein 2 (*CAVIN2*), Dynactin Subunit 4 (*DCTN4*), Eukaryotic Translation Termination Factor 1 (*ETF1*), *GIMAP4-201* transcript of the gene GTPase, IMAP Family Member 4, and *NUDT21-201* transcript of the gene Nudix Hydrolase 21 (*NUDT21*) (Fig.4a). The IR node *NUDT21-201* appears as a master hub of degree 17 in the affected individual’s network and is additionally a DE transcript (BH, p = 0.002, logFC = −0.11). The corresponding *NUDT21* gene is a known splicing factor and was identified in a recent study as a differentially expressed RNA-modification-related gene in MDD ^74^. In Table S11, we provide a list of all 61 master hub nodes in the AIN, with 30 (49%) showing evidence for association with psychiatric disorders, 16 of which appear in our analysis at the transcript level as IR master hubs.

**Figure 4:**
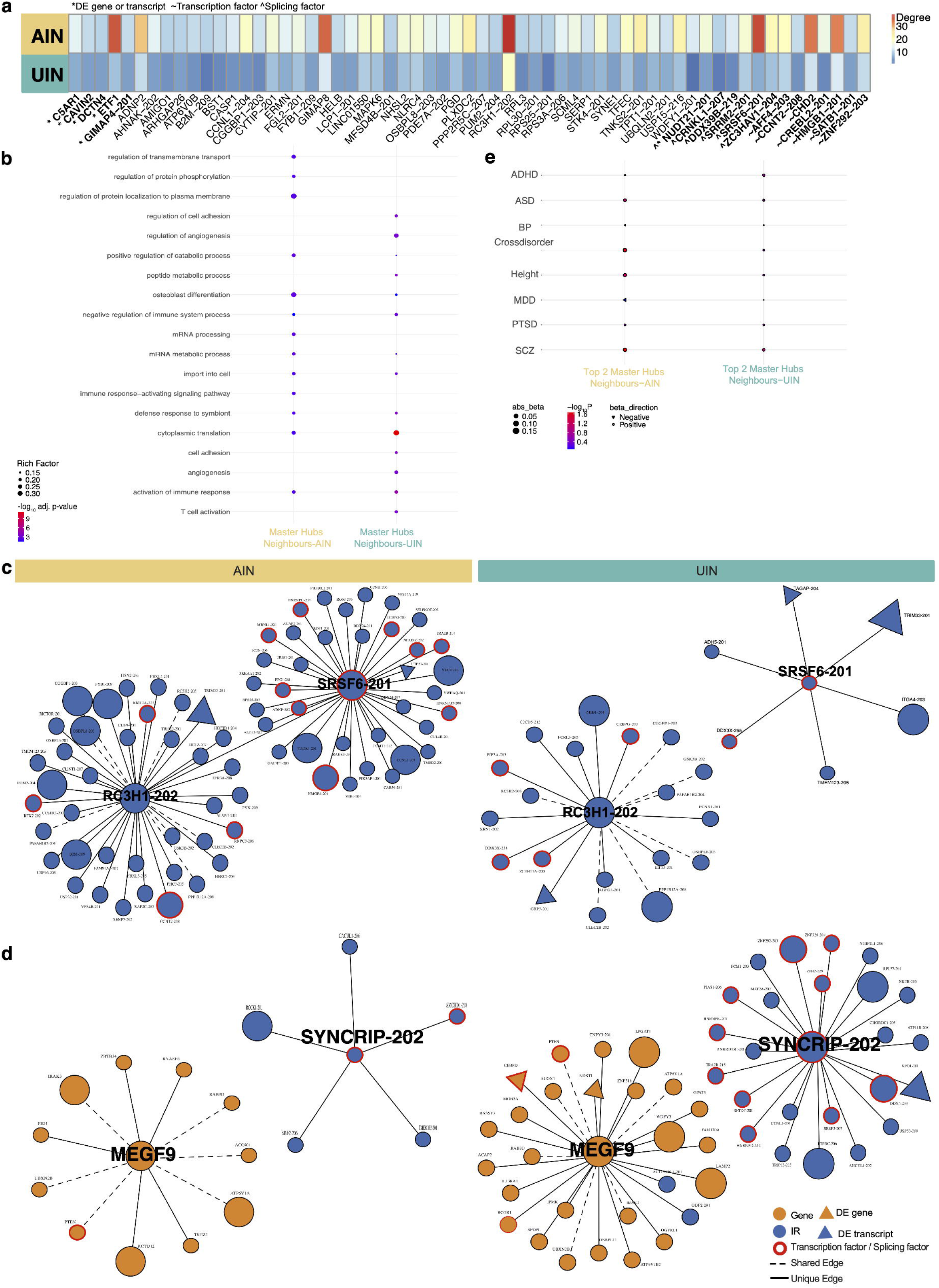
**a** Heatmap of the 61 master hub nodes in the AIN whose degree increased by more than two-fold compared to the un-thresholded UIN. Among the 61 master hub nodes are six differentially expressed genes and transcripts (*), seven TFs (~), and six SFs (^). **b** Enrichment in GO biological processes showing the top 10 (based on BH adj p-values) enriched processes for the 61 master hubs in each network and their first-order neighbors. **c** Top 2 master hubs (highest degree) in the AIN (left) showing distinct connection patterns compared to the corresponding nodes in the UIN (right). **d** Top 2 master hubs in the UIN (right) showing distinct connection patterns compared to the corresponding nodes in the AIN (left). **e** Enrichment of the top 2 master hubs in each network and their two-hop neighbors in genes that carry SNPs associated with the GWAS of different psychiatric disorder traits and Height.

Similarly, we found 61 master hubs in the UIN showing twice as much connectivity compared to the AIN. A GO enrichment analysis of each network’s master hubs (61 each) and their first-order neighbors, showed distinct biological processes related to each network (Fig.4b, Table S12). The set of master hubs of the UIN and their first-order neighbors indicated enrichments in cytoplasmic translation, activation of immune response, and cell adhesion processes (BH, p < 0.05), while the set of master hubs of the AIN and their first-order neighbors showed enrichment in other biological processes, including positive regulation of catabolic process, osteoblast differentiation, and mRNA processes.

To further investigate whether master hubs reflect network-specific biology, we focused on the top two master hubs with the highest degree in each network. Figure 4c illustrates the top two master hubs in the AIN, *RC3H1-202* transcript of the gene Ring Finger And CCCH-Type Domains 1, and *SRSF6-201* transcript of the gene Serine and Arginine Rich Splicing Factor 6, along with their corresponding counterparts in the UIN. Similarly, Figure 4d presents the top two master hubs in the UIN, the Multiple EGF Like Domains 9 gene (*MEGF9*), and *SYNCRIP-202* transcript of the gene Synaptotagmin Binding Cytoplasmic RNA Interacting Protein (*SYNCRIP*), along with their respective versions in the AIN. Focusing on these top two master hubs and their two-hop neighbors, we conducted a GWAS enrichment analysis for each network. Specifically, we investigated enrichment in genes harboring SNPs associated with psychiatric disorders (see Methods section 8.1). Our analysis revealed significant enrichment of *RC3H1-202*, *SRSF6-201*, and their two-hop neighbors (328 genes) in the AIN for genes implicated in cross disorder and schizophrenia (β = 0.18, p = 0.03, β = 0.16, p= 0.02 respectively). No corresponding significant enrichment was found using the set of genes comprising the master hubs *MEGF9*, *SYNCRIP-202*, and their two-hop neighbors (259 genes) in the UIN, see Fig.4e and Table S13.

#### 3.3. Isoform ratio nodes show distinct co-regulatory patterns compared to their total expression nodes and capture different protein-protein interactions

Our analysis found that hub nodes are enriched for IR nodes, with the proportion of IR nodes exhibiting a positive correlation with increasing hub degree (e.g., ≥ 15 and ≥ 20, see Fig.5a). This observation underscores the critical role of IR nodes within these networks. To highlight the importance of IR nodes, we present as an example *HNRNPH1-227*, a hub IR node (degree ≥ 10), and a known transcription and splicing factor associated with neurodevelopmental disorders ^75,76^. This node exhibits distinct connection patterns compared to its corresponding TE gene node in both networks (Fig.5b,c). Investigation of this node in the BioGrid database ^77^ showed that, at the isoform level, *HNRNPH1-227* establishes connections with a unique set of proteins, diverging from those observed at the gene level (Fig.5d). More specifically, within the AIN, *HNRNPH1-227* interacts with isoforms of the proteins RPS6, and EIF4B, whereas the corresponding TE gene node captures interactions with EWSR1, SF1, HNRNPH3, HNRNPA1, and FUS. Likewise, within the UIN, *HNRNPH1-227* interacts with isoforms of the proteins CHD2, and DDX17, while the corresponding TE gene node interacts with EWSR1, HNRNPA1, SRSF5, HNRNPA2B, and ILF3. The observed differences in interaction patterns between isoform-specific nodes and their gene-level counterparts highlight the potential for isoform-level analysis to reveal previously unrecognized molecular relationships and functional specificities.

**Figure 5:**
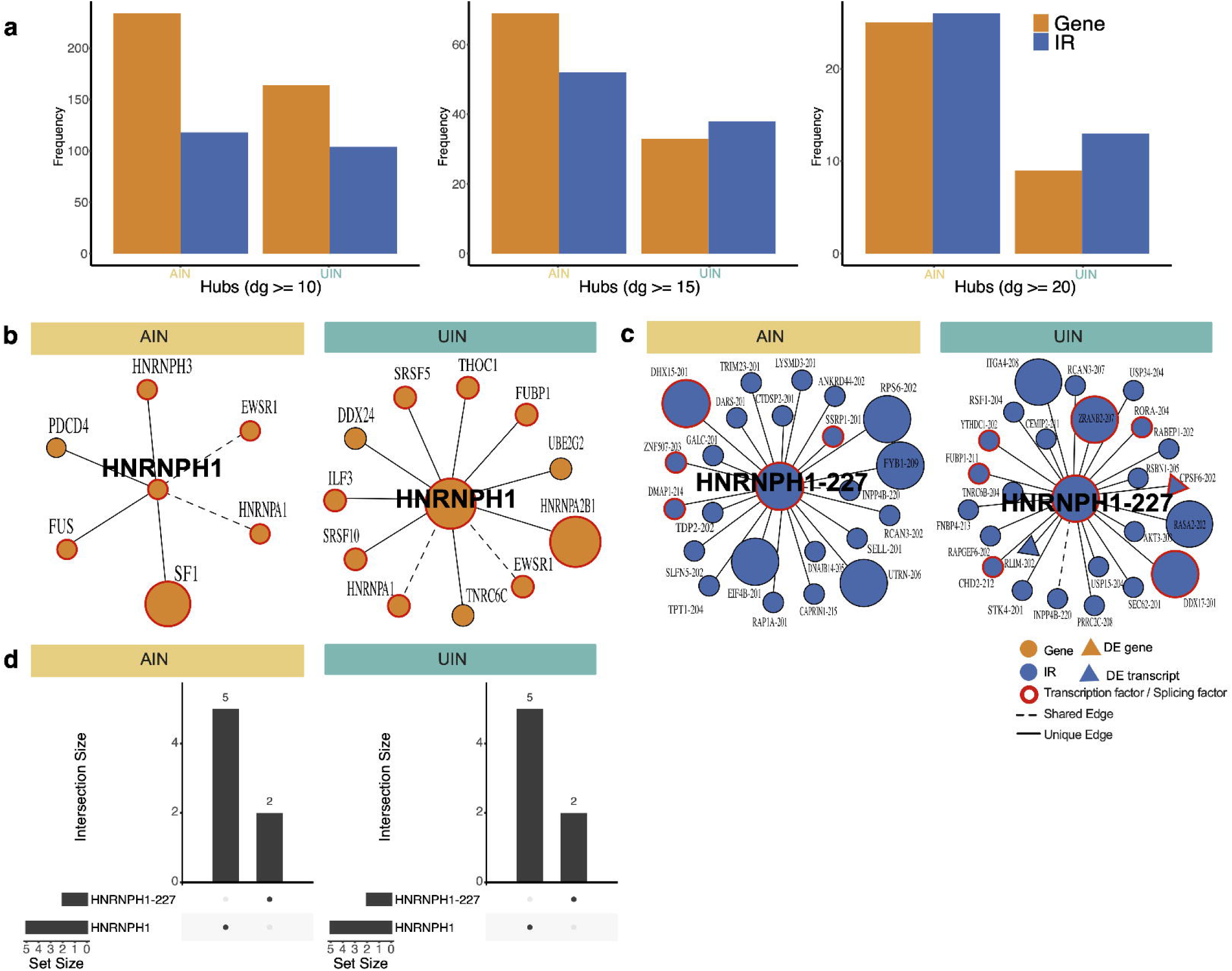
**a** IR versus TE nodes distribution in hubs of degree >=10 (left), degree >=15 (middle), and degree >= 20 (right) in both the affected and unaffected individuals’ networks. **b** First-order neighbors of the transcription and splicing factor gene *HNRNPH1* in the affected and unaffected individuals’ networks. *HNRNPH1* is known to be associated with neurodevelopmental disorders and shows distinct connection patterns in each network. **c** The hub IR node *HNRNPH1-227* shows distinct first-order neighbors in each network that also differ from the neighbors of its TE node *HNRNPH1*. **d** Upset plot showing that a distinct set of protein-protein interactions are captured at gene (*HNRNPH1*) vs isoform-level (*HNRNPH1-227*) in both networks.

## Discussion

In this work, we present an attempt to integrate gene expression and isoform ratio data into a comprehensive network approach to investigate co-expression patterns in individuals with and without stress-related psychiatric disorders, with a focus on depressive symptoms. This integrative approach allowed us to capture a deeper understanding of the complex interplay between multiple genes and isoforms, rather than focusing on the traditional gene-level analysis, revealing significant differences in network architecture between the two groups of individuals. We identified a total of 450 differentially expressed genes and 269 differentially expressed transcripts, including 104 with isoform-specific differential expression. Enrichment analyses further highlighted the unique signal arising from isoform-level data to understand gene regulation in stress-related psychiatric disorders.

While differential expression analysis has been widely used to study psychiatric disorders ^6,8–10^, incorporating isoform-level data into such analyses remains a relatively unexplored area. We compared our results to studies using blood samples from patients with MDD. Ota et al. ^9^ found 8 differentially expressed genes in a longitudinal study of children and adolescents, with one gene overlapping with our DE genes: NADH:Ubiquinone Oxidoreductase Subunit A2 (*NDUFA2*). Wittenberg et al. ^10^ conducted a meta-analysis of MDD across several blood transcriptome studies and provided a harmonized list of 272 DE genes. Of these, only six overlapped with our DE genes with concordant direction, namely *MBNL1* Antisense RNA 1 (*MBNL1-AS1*), IKAROS Family Zinc Finger 3 (*IKZF3*), and BUB3 mitotic checkpoint protein (*BUB3*) as down-regulated, and WD repeat domain 74 (*WDR74*), N-deacetylase and N-sulfotransferase 1 (*NDST1*), and Fc alpha receptor (*FCAR*) as up-regulated. Moreover, from the harmonized gene list, two genes, namely calcium/calmodulin-dependent protein kinase kinase 2 (*CAMKK2*), and protein tyrosine phosphatase non-receptor type 4 (*PTPN4*) appear in our analysis as DE solely at the transcript level (*CAMKK2-209*, and *PTPN4-201* respectively) with no corresponding gene-level DE. Our DTA and the unique enrichment profile of differentially expressed transcripts highlight the importance of isoform-level data, as it captures additional layers of transcriptional complexity that may be missed by traditional gene-level analyses.

While DE analysis provides information about individual genes and transcripts, it examines them in isolation, failing to capture the complex interplay between multiple genes and transcripts. Network co-expression analysis approaches represent a valuable tool, providing a comprehensive framework that captures the intricate interactions and functional relationships between genes and transcripts. Many studies have shown that the wiring of networks changes under different conditions, reflecting the pathophysiological states associated with diseases ^78–81^, such as breast cancer ^80^. In psychiatric disorders, existing network approaches have focused on DE genes as input for network inference ^15,82,83^. However, hub nodes in biological networks have been found to play a critical biological function even if the gene itself does not show differential expression ^7,80,84,85^. Moreover, most methods focused on gene-level networks without considering isoform-level co-regulatory changes ^8,11,12,82,83^. While gene-level analyses provide valuable insights, incorporating isoform-level data is crucial for capturing the full complexity of gene regulation in psychiatric disorders. Isoforms can have distinct functions and interactions, and their dysregulation may contribute to disease pathogenesis in ways that are not apparent at the gene level^24,25,86^.

Therefore, we introduce an integrative approach combining both total gene expression values (TEs) and isoform ratios (IRs) to construct separate co-expression networks for affected and unaffected individuals (AIN vs. UIN), thereby using all genes and transcripts that passed our quality control (n=7394, n=7097). We ensured the robustness of ARACNE network inference in capturing biological differences between the two groups using advanced graph generation and embedding techniques. This marks the first attempt to leverage AI techniques, such as graph embeddings and graph generation, for the validation of network inference, enhancing the reliability of the network analysis results by confirming the distinctiveness of the network topologies. Hub nodes were a key focus of our network analysis. We identified hub nodes in both networks, with 127 common hubs showing distinct connection patterns between the AIN and UIN. Specifically, 82.4% of the edges connected to these common hubs in the AIN were unique to that network, while 79.7% of the edges connected to the same hubs in the UIN were unique to the UIN. This suggests a potential rewiring of regulatory interactions in psychiatric disorders, which could have downstream effects on pathways and cellular functions, such as synaptic plasticity, neurotransmitter signaling, or the immune response. Further investigation of these altered hub connections may provide insights into the molecular mechanisms underlying disease pathogenesis and identify potential therapeutic targets.

Focusing on the AIN, our analysis revealed 61 master hub nodes, with over half being isoform ratio nodes. Master hubs represent highly influential nodes, exhibiting at least two-fold degree change in one network compared to the other. The fact that over half of these master hubs were isoform ratio nodes underscores the potential of isoform-specific dysregulation to drive network alterations in affected individuals. Investigation of these master hubs in the AIN showed significant associations with psychiatric disorders, particularly those related to psychiatric cross-disorder phenotype, suggesting their involvement in shared disease processes across different diagnoses. In a study by Wei et al.^83^ investigating MDD in the dentate gyrus (DG) and anterior cingulate cortex (ACC) regions of a mouse brain, the authors created an interaction network of DE genes and identified important differentially expressed TFs that regulate many hubs, including the TF gene chromodomain helicase DNA binding protein 2 (*CHD2*). *CHD2* was predicted to upregulate the expression of DE genes related to MDD in the DG region. In our analysis, *CHD2* appears as an important gene with differential expression found at the transcript level (*CHD2-201*). This transcript-level DE provides a more nuanced view of *CHD2* dysregulation compared to previous studies that focused on gene-level expression. Furthermore, *CHD2* is a master hub node in the AIN with a degree of 34 compared to 12 in the UIN, suggesting altered regulatory interactions in the context of psychiatric disorders. Further investigation of the specific functions and interactions of *CHD2* isoforms may reveal novel therapeutic targets for psychiatric disorders, particularly depression which represents the major portion of the clinical phenotype studied here.

To further investigate the functional roles of master hubs, we performed pathway enrichment analysis of the master hubs in each network and their first-order neighbors. The set of master hubs of the AIN and their first-order neighbors showed enrichment in positive regulation of catabolic processes, osteoblast differentiation, and mRNA processes. The enrichment of mRNA processes in the AIN could suggest dysregulation of RNA processing and splicing, potentially leading to altered isoform expression and downstream functional consequences. While seemingly unrelated to psychiatric disorders at first glance, enrichment in osteoblast differentiation may have intriguing implications. Osteoblast differentiation is crucial for bone formation and remodeling, and dysregulation of this process is implicated in osteoporosis. Studies have shown a complex interplay between osteoporosis, chronic stress, and inflammation ^87^. Chronic stress, a well-established risk factor for psychiatric disorders like depression, can trigger increased inflammatory factors that influence osteoblast differentiation and contribute to osteoporosis. This suggests a potential link between the enriched osteoblast differentiation pathway in the AIN and the chronic stress often experienced by individuals with psychiatric disorders. Further investigation of this pathway may uncover novel connections between bone health, stress response, and mental health.

Since the UIN represents the healthy control group, pathways enrichments of the master hubs and their first-order neighbors may reflect normal biological processes and regulatory pathways that are essential for maintaining healthy functions in the body. For instance, the enrichment of cytoplasmic translation processes could be crucial for supporting protein synthesis and synaptic plasticity, which are essential for learning, memory, and cognitive flexibility. Since the list of significant biological processes associated with master hubs in the AIN and UIN is fully distinct, it suggests that the two networks represent different molecular pathways involved in disease pathogenesis, potentially leading to the identification of network-specific therapeutic targets.

Our analysis found that hub nodes were enriched in isoform ratio nodes which captured unique protein-protein interactions compared to their corresponding TE nodes. For example, the IR node *HNRNPH1-227* exhibited distinct protein-protein interactions compared to its gene-level node. This observation highlights the potential for isoform-level analysis to reveal previously unrecognized molecular relationships and functional specificities. We also investigated our network properties at the domain level. Domains are distinct functional and structural units within proteins that mediate interactions with other molecules. Using the domain interaction exploratory tool DIGGER ^71^, we found evidence suggesting domain-level changes, highlighting the importance of a higher-level understanding of system interactions.

Throughout our analysis, we highlighted examples of hub and master hub nodes associated with psychiatric disorders, providing concrete evidence for the relevance of our network-based approach. However, this work also presents opportunities for future research to build upon our findings. For example, while we used ARACNE for network inference, which effectively captures non-linear relationships between genes and isoform ratios, future studies could assess the robustness of our findings using alternative approaches. Similarly, while our network analysis reveals co-regulatory changes and network-specific biomarkers, exploring alternative methods for network differential analysis like BoostDiff ^88^ and chNet ^80^, could provide an alternative way to understand network alterations. Additionally, exploring gene expression patterns in other tissues, such as the central nervous system or the gut, could provide a more comprehensive understanding of the molecular processes underlying psychiatric disorders ^89^. Further, investigating the potential influence of pharmacotherapy and analyzing longitudinal data could reveal additional insights into disease mechanisms and progression. The cell type heterogeneity in blood samples, especially the imbalance between PBMCs used for affected individuals and the mix of PBMCs and whole blood samples for unaffected individuals, could have affected our results. While we adjusted for different blood cell type compositions, further investigation using more homogenous cell populations or advanced deconvolution methods could refine our understanding of cell-type-specific effects. Finally, exploring alternative representations of splicing events, such as exon-level values, could offer more granular insights into splicing dynamics.

Despite these limitations, our study provides novel evidence for the importance of isoform-level analysis in understanding the complex landscape of gene regulation in psychiatric disorders. Consequently, our findings emphasize the need for comprehensive functional annotation of isoforms to better understand their roles in complex biological processes and disease mechanisms.

## Data and Code Availability

The code is available on https://github.com/cellmapslab/NetIso/, and the RNA-seq data on GEO under accession numbers GSE289144 and GSE289146.

## Acknowledgments

We extend our gratitude to all participants of the BeCOME and OPTIMA studies without whom our research would not be possible. Markus List’s contributions were supported by the Klaus Tschira Stiftung (KTS, Klaus Tschira Foundation, Grant 00.003.2024). Annalisa Marsico and Lambert Moyon acknowledge support by the BMBF Cluster4Future program CNATM. Dr. Knauer-Arloth’s contributions were supported by the Brain & Behavior Research Foundation (NARSAD Young Investigator Grant, #28063). Ghalia Rehawi is supported by the Helmholtz Association under the joint research school “Munich School for Data Science - MUDS”.

## CRediT authorship contribution statement

**Ghalia Rehawi**: Writing – original draft, Visualization, Software, Methodology, Formal analysis, Conceptualization. **Jonas Hagenberg**: Software, review & editing. **Philipp Sämann**: Data curation, review & editing. **Lambert Moyon**: review & editing. **Elisabeth Binder**: Funding acquisition. **Annalisa Marsico**: Conceptualization, Supervision, review & editing. **Markus List**: Supervision, review & editing. **Janine Knauer-Arloth**: review & editing, Visualization, Supervision, Conceptualization, Funding acquisition.

