## Supplementary Materials for "Integrative Gene and Isoform Co-expression Networks Reveal Regulatory Rewiring in Stress-related Psychiatric Disorders"

### 1. Supplementary Methods

#### 1.1. Cell type deconvolution

Gene and transcript-level reads were filtered for unwanted sequences using Cutadapt ^1^ v2.10, we also removed zero-length reads and retained only those with a count ≥ 10 in at least 95% of samples, resulting in 9777 genes and 11427 transcripts. Before correcting for batch effects, we calculated the cell type decomposition values using the Granulator v1.2.0 ^2^ package on the raw gene counts of the 9777 genes. We used the leukocyte gene signature matrix LM22 ^3^ as a reference matrix and applied the dtangle algorithm from Granulator resulting in 22 cell types: B cells memory, B cells naive, Dendritic cells activated, Dendritic cells resting, Eosinophils, Macrophages M0, Macrophages M1, Macrophages M2, Mast cells activated, Mast cells resting, Monocytes, Neutrophils, NK cells activated, NK cells resting, Plasma cells, T cells CD4 memory activated, T cells CD4 memory resting, T cells CD4 naive, T cells CD8, T cells follicular helper, T cells gamma delta, T cells regulatory (Tregs). Principal components (PCs) of the cell type proportions for the 336 individuals (107 unaffected and 229 affected) were calculated, and the first 5 PCs covering 54% of the variance were included for the batch correction process.

#### 1.2. Batch correction

To account for confounding effects, we first corrected the gene and transcript-level data for the sequencing run using the `removeBatchEffect` function from the limma R package V3.50.1 ^4,5^. The sequencing run constituted a huge effect which can be seen in a PC analysis in Figure S1A. We then performed Surrogate Variable Analysis (SVA) ^6^ and identified 21 additional hidden batch effects. The decision on which variables to include for batch correction was carried out in two steps: 1) a canonical correlation analysis which identified correlations between the known biological and technical covariates (sex, age, BMI, the 5 PCs of cell type proportions, GC content, total read pairs, study, status, RNA integrity number (RIN), study, plates, and preparation batch), the 21 SVs, and the first 10 PCs of the gene counts after correcting for sequencing run (Figure S1B). Most importantly, it identified a correlation between PC1, SV1, GC content, total read pairs, and RIN. It also identified a correlation between the second principal component of cell type decomposition values (cellType_pc_2) and status. 2) association analysis using ANOVA of the aforementioned technical and biological covariates, top 5 SVs extracted after correcting for sequencing run with the first 10 PCs of the gene counts after correcting for sequencing run (Figure S1C). The results showed that SV1, GC content, total read pairs, and RIN are significantly associated with PC1. Hence the final set of variables we correct for includes GC content, total read pair, and the top 5 PCs of cell type proportions. We disregard RIN to avoid overcorrection since it correlated with total read pairs. Although not shown to be significant contributors to variance in the data, we nonetheless correct for sex, age, and BMI as those variables have been shown in previous studies to be important confounders. We removed genes and transcripts that had negative values due to the subtraction of the modeled batch effects resulting in 7394 genes and 7334 transcripts.

#### 1.3. ARACNE for network inference

For both the affected individuals (AIN) and unaffected individuals' networks (UIN), we used ARACNE (Algorithm for the Reconstruction of Accurate Cellular Networks) ^7^, an information theoretic-based method designed for the reverse engineering of regulatory networks. Specifically, ARACNE builds a network in two steps. In the first step, the mutual information between each pair of input values is calculated and treated as edge weight. In the second step, the majority of false positive interactions are removed by applying the Data Processing Inequality (DPI) ^8^, where an edge with the smallest MI value in all network triplets is removed and regarded as an indirect interaction. For example, the weakest edge in a triplet (i,j,k), say (i,k), is removed if $MI(X_{ik}) \leq min(MI(X_{ij}), MI(X_{jk}))$.

### 2. Supplementary Results

#### 2.1. Sample size effect analysis

#### In an initial analysis, our samples comprised affected and unaffected individuals only from the OPTIMA and BeCOME cohorts, with a total number of 210 affected and 63 unaffected individuals after outlier filtering. This huge sample size difference affects the network inference step, leading to uncertain conclusions regarding differences in the networks’ structures. Figure S3 shows a PCA of the embeddings of the AIN (n=210), the UIN (n=63), as well as 10 different sub-AINs constructed by randomly sampling 60 samples, and 10 sub-AINs constructed by randomly sampling 150 samples from the affected group. These embeddings were calculated using the Graph2Vec technique ^9^, with an embedding vector dimension of 128. The results show the effect of sample size on the inferred networks’ structures which can be seen at the embedding level and is captured by PC1. Specifically, sub-AINs constructed with a higher sample size (150) are closer in the embedding space to the affected individuals’ network constructed using the entire available samples (210) compared to the sub-AINs which are built using a smaller sample size (60). Therefore, to guarantee a high-quality network inference, unbiased by sample size effects, and enabling fair comparison of the two networks, we increased the sample size of the unaffected group by integrating the IST study, adding 37 control individuals. We would like to highlight that this analysis investigating the effect of sample size on network inference using graph embedding approaches is a novel effort that has not been explored before in the literature.

#### 2.2. Network inference for affected and unaffected individuals

We infer the AIN for the affected group (n=210) and the UIN for the unaffected group (n=95) using both total gene expression (n=7394) and isoform ratios (n=7097) as input for ARACNE. Following the work of ^10^, we filter edges connecting features of the same gene to reduce bias in the networks and increase interpretability. This means we remove edges of type IR-IR connecting two isoforms of the same gene and remove edges of type TE-IR connecting a gene with its isoform. The resulting AIN consists of 14,300 nodes and 21,324 edges, whereas the UIN consists of 14,447 nodes and 40,676 edges. This difference in the number of inferred edges between the two networks is due to the heterogeneity of the sample size since the UIN is built using a smaller sample size. This causes the UIN to have a higher number of edges with lower MI values compared to the AIN (Figure S4B). Hence, to ensure a fair comparison of the two networks we need to balance the number of edges and ensure that only edges with high MI values are included in the final networks. To this end, we filter out edges with low MI values in both networks. For each edge type, we choose one threshold as the edge type-specific median MI value in the AIN, since it is more robustly constructed with a higher number of samples i.e. 0.24 for TE-TE edge type, 0.14 for TE-IR edge type, and 0.38 for IR-IR edge type (Figure S4A). We use this filtering threshold for both networks to obtain the final filtered networks (Figure S4C-D). This thresholding leads to the two networks having similar numbers in terms of #nodes, #edges, and #edges per edge type (Table 2), guaranteeing a fair comparison between the two networks.

### Supplementary Figures

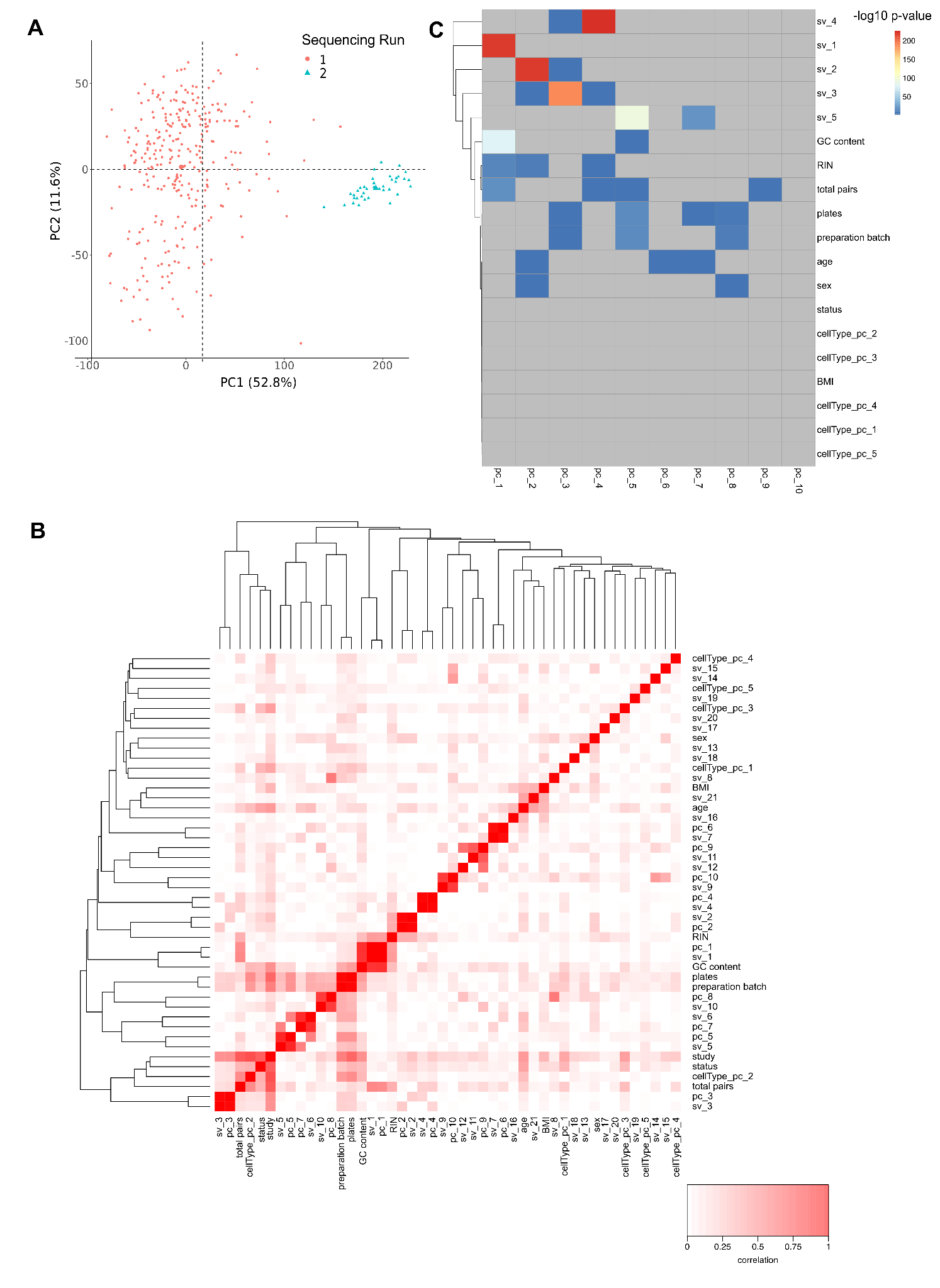

**Figure S1**: (A) The first and second principal components of a PCA show the separate distribution of samples based on the RNA-sequencing run, 1) samples sequenced from OPTIMA and BecOME cohorts and 2) samples sequenced separately from the IST study. (B) Heatmap showing the canonical correlations between all pairwise variables including technical and biological covariates, 21 SVs extracted after correcting for sequencing run, the first 5 PCs of cell type decomposition values, and the first 10 PCs of the gene counts after correcting for sequencing run. (C) Heatmap of -log_10_ p values from ANOVA tests between technical and biological covariates, 5 SVs extracted after correcting for sequencing run, the first 5 PCs of cell type decomposition values, and the first 10 PCs of the gene counts after correcting for sequencing run.

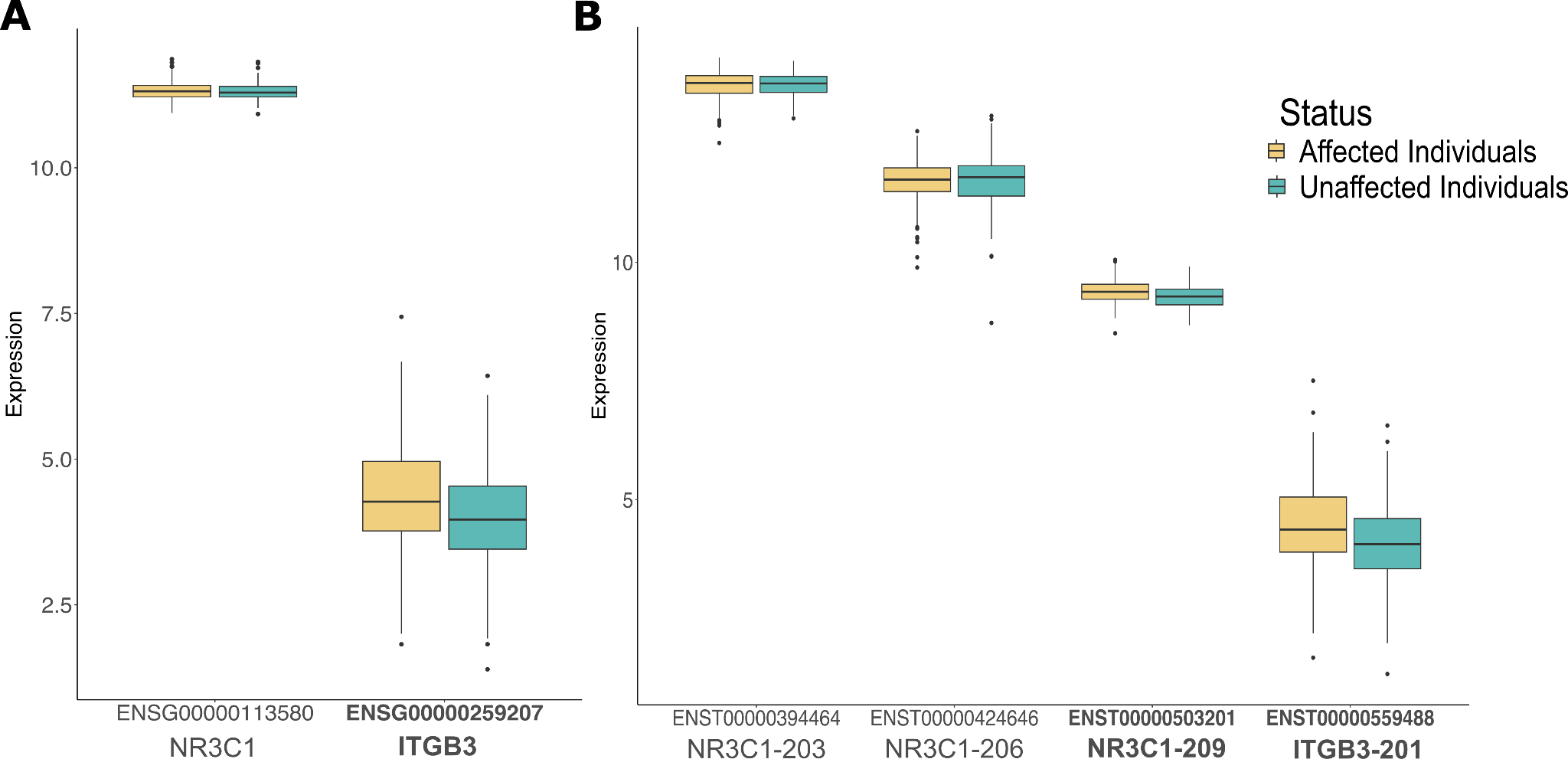

**Figure S2**: (A) The expression in the affected and the unaffected groups of two genes *ITGB3* (ENSG00000259207) which is up-regulated (bold) and *NR3C1* (ENSG00000113580) which is not differentially expressed. (B) The expression in the affected and the unaffected groups of 4 transcripts, 3 transcripts of the gene *NR3C1*, which does not show differential expression at the gene level, but one of its transcripts *NR3C1-209* (ENST00000503201) is up-regulated (bold). The up-regulation of the gene *ITGB3* is also visible at its transcript level where *ITGB3-201* (ENST00000559488) is also up-regulated (bold).

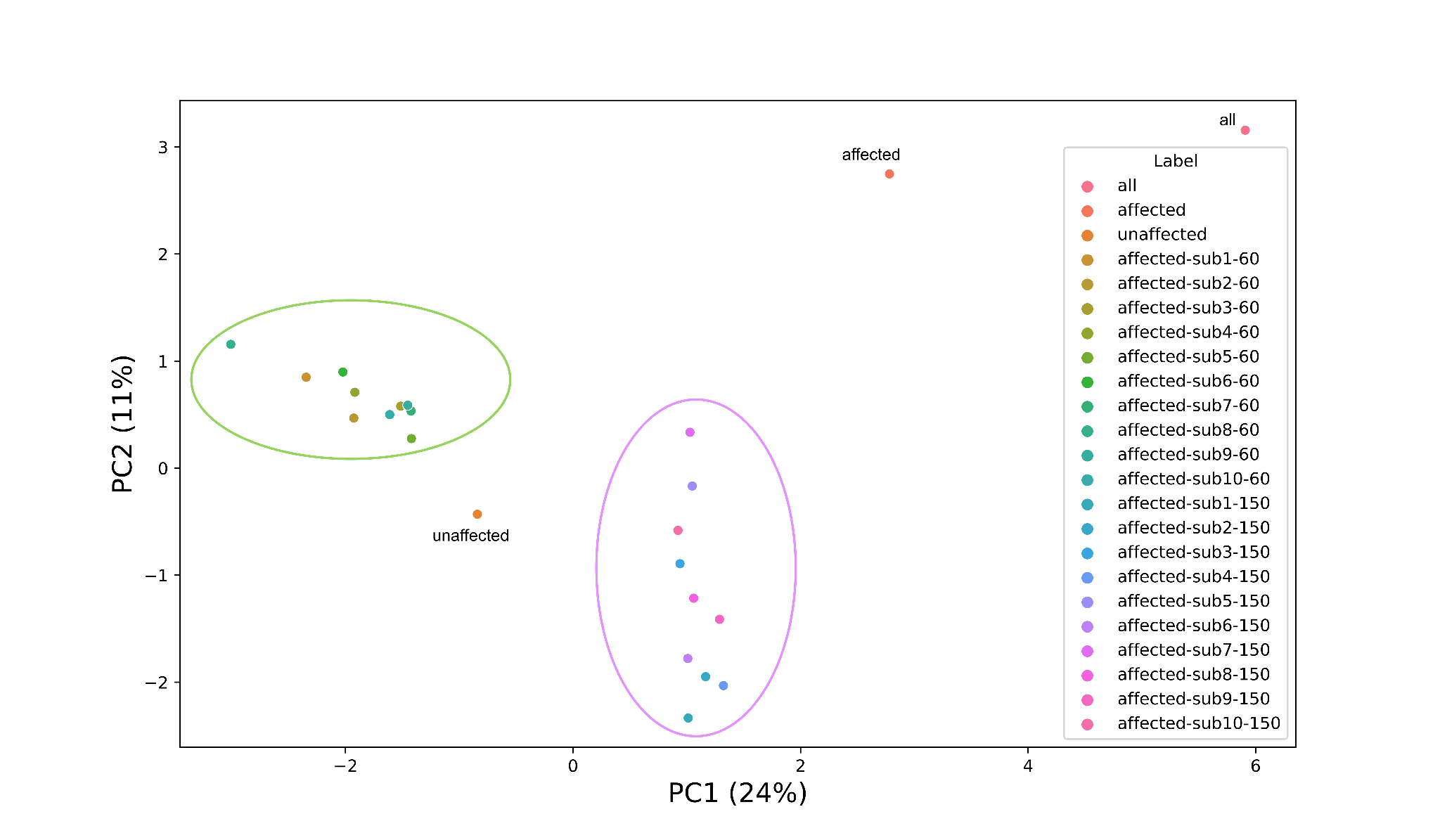

**Figure S3**: The first and second PCs of a PCA on graph embeddings obtained using Graph2Vec. The graph embeddings are of networks of different sizes: the full affected individuals’ network (n=210), the full unaffected individuals’ network (n=63), a combined network for affected and unaffected individuals (all), 10 sub-networks built using 60 samples from the affected group (green ellipse), and 10 sub-networks built using 150 samples from the affected group (violet ellipse).

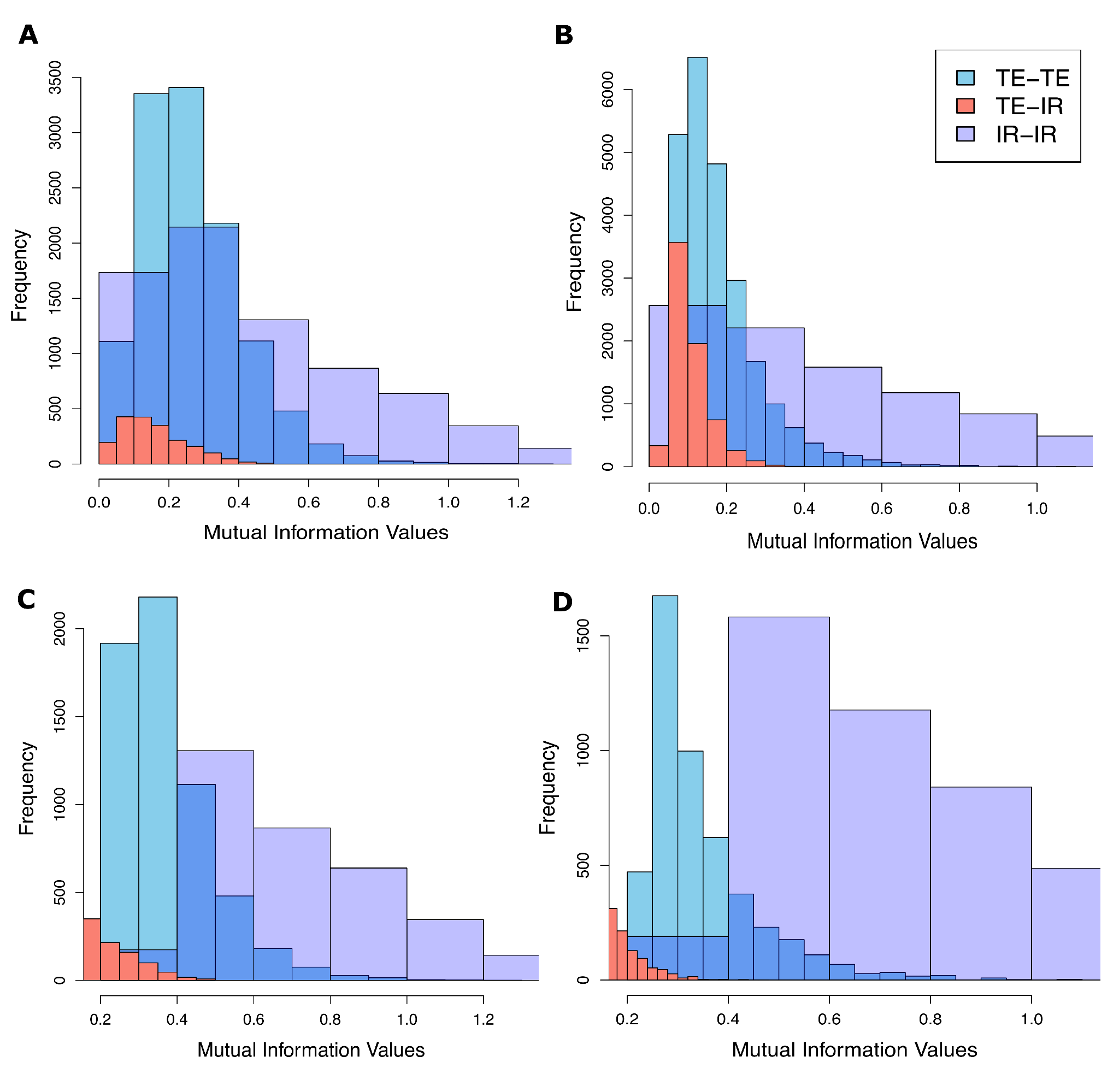

**Figure S4**: (A) The MI distribution of edges in the AIN before thresholding: Median= 0.24, Mean= 0.26 for TE-TE edge types, Median= 0.14, Mean= 0.156 for TE-IR edge types, and Median= 0.38, Mean= 0.49 for IR-IR edge types. (B) The MI distribution in the UIN before thresholding: Median= 0.15, Mean= 0.18 for TE-TE edge types, Median= 0.09, Mean= 0.10 for TE-IR edge types, and Median= 0.39, Mean= 0.50 for IR-IR edge type. (C) The MI distribution in the AIN after thresholding: Median= 0.34, Mean= 0.37 for TE-TE edge types, Median= 0.21, Mean= 0.23 for TE-IR edge types, and Median= 0.68, Mean= 0.77 for IR-IR edge types. (D) The MI distribution in the UIN after thresholding: Median= 0.31, Mean= 0.35 for TE-TE edge types, Median= 0.17, Mean= 0.19 for TE-IR edge types, and Median= 0.71, Mean= 0.81 for IR-IR edge types.

### Supplementary Tables

**Table S1:** This table contains gene-disease associations (GDAs) for various psychiatric disorders extracted from the DisGeNET database. The table includes the following columns: official gene symbol (Gene), full name of the gene (GeneFullName), name of the psychiatric disorder (Disease), DisGeNET score for the gene-disease association (values > 0.4) (ScoreGDA), number of PubMed IDs supporting the association (NumPMIDs), and number of SNPs associated with the gene (NumVariantsAssociatedToGene)**.**

**Table** **S2:** This table provides a comprehensive list of known transcription factors and splicing regulators compiled from multiple resources. It includes the following columns: the official gene symbol (gene_symbol), the type of the regulatory gene, and whether it is a transcription factor (TF) or splicing factor (SF) (type).

**Table S3:** This table presents our differential gene expression analysis results comparing individuals affected by psychiatric disorders to unaffected controls. The table includes the following columns: ensemble gene IDs (gene_id), log2 fold change in expression (affected vs. unaffected) (logFC), average expression level across all samples (AveExpr), t-statistic from the differential expression analysis (t), unadjusted p-value (P.Value), adjusted p-value (FDR) (adj.P.Val), log-odds that the gene is differentially expressed (B), official gene name (gene_name).

**Table S4:** This table presents our differential transcript expression analysis results comparing individuals affected by psychiatric disorders to unaffected controls. The table includes the following columns: ensemble transcript IDs (transcript_id), log2 fold change in expression (affected vs. unaffected) (logFC), average expression level across all samples (AveExpr), t-statistic from the differential expression analysis (t), unadjusted p-value (P.Value), adjusted p-value (FDR) (adj.P.Val), log-odds that the gene is differentially expressed (B), official transcript name (transcript_name).

**Table S5:** This table provides the results of Gene Ontology (GO) enrichment analysis for differentially expressed genes and differentially expressed transcripts. The table includes the following columns: GO term identifier (ID), description of the GO term (Description), ratio of genes in the input list associated with the GO term (GeneRatio), ratio of background genes associated with the GO term (BgRatio), the ratio of input genes that are annotated in a term to all genes that are annotated in this term (RichFactor), the ratio of the frequency of input genes annotated to a particular GO term compared to the frequency of all genes annotated to that term (foldEnrichment), statistical significance measure of the enrichment (zScore), unadjusted p-value for enrichment (pvalue), adjusted p-value (BH) (p.adjust), Q-value for enrichment (qvalue), list of genes associated with the GO term (geneID), number of genes in the input list associated with the GO term (count), the set on which enrichment analysis is being done (set).

**Table S6:** This table presents the results of MAGMA gene-set analysis, assessing the enrichment of DE genes and DE transcripts in genes carrying single nucleotide polymorphisms (SNPs) associated with psychiatric disorders. The table includes the following columns: input gene list used for enrichment (VARIABLE), name of the psychiatric disorder or trait analyzed (TYPE), number of genes in the set (NGENES), effect size (positive values indicate enrichment) (BETA), standardized effect size (BETA_STD), standard error of the effect size (SE), p-value for enrichment (P), absolute value of the effect size beta (abs_beta), direction of the effect (beta_direction), adjusted p-value (BH) (adjP_val), -log10 of adjusted p-value ((minus) log10_adj_p).

**Table S7:** This table contains gene-disease associations based on the intersection of all DEGs and DETs with genes of Table S1, resulting in 53 genes. The table includes the following columns: official gene symbol (Gene), full name of the gene (GeneFullName), name of the psychiatric disorder (Disease), DisGeNET score for the gene-disease association (values > 0.4) (ScoreGDA), number of PubMed IDs supporting the association (NumPMIDs), and number of SNPs associated with the gene (NumVariantsAssociatedToGene)**.**

**Table S8:** This table contains information about the 352 hubs (degree >= 10) in the affected individuals’ network. The table includes the following columns: gene and transcript symbols of TE and IR hub nodes (node), the degree of the node (degree), and ensemble IDs (id).

**Table S9:** This table contains information about the 268 hubs (degree >= 10) in the unaffected individuals’ network. The table includes the following columns: gene and transcript symbols of TE and IR hub nodes (node), the degree of the node (degree), and ensemble IDs (id).

**Table S10:** This table contains the common hub nodes with degree >= 10 in both networks. The table includes the following columns: gene and transcript symbols of TE and IR hub nodes (hubs_intersect).

**Table S11:** This table contains a list of all 61 master hub nodes in the AIN, with 30 (49%) showing evidence for association with psychiatric disorders based on a manual search of PubMed. The table includes the following columns: gene or transcript official symbol (Master_hub_node_AIN), link to PubMed publication showing evidence of association with psychiatric disorders (Evidence1), and further evidence (Evidence2).

**Table S12:** This table provides the results of Gene Ontology (GO) enrichment analysis for the 61 master hubs in each network and their first-hob neighbors. The table includes the following columns: GO term identifier (ID), description of the GO term (Description), ratio of genes in the input list associated with the GO term (GeneRatio), ratio of background genes associated with the GO term (BgRatio), the ratio of input genes that are annotated in a term to all genes that are annotated in this term (RichFactor), the ratio of the frequency of input genes annotated to a particular GO term compared to the frequency of all genes annotated to that term (foldEnrichment), statistical significance measure of the enrichment (zScore), unadjusted p-value for enrichment (pvalue), adjusted p-value (BH) (p.adjust), Q-value for enrichment (qvalue), list of genes associated with the GO term (geneID), number of genes in the input list associated with the GO term (count), the set on which enrichment analysis is being done (set).

**Table S13:** This table presents the results of MAGMA gene-set analysis, assessing the enrichment of the top two master hubs with the highest degree in each network and their two-hob neighbors in genes carrying single nucleotide polymorphisms (SNPs) associated with psychiatric disorders. The table includes the following columns: input gene list used for enrichment (VARIABLE), name of the psychiatric disorder or trait analyzed (TYPE), number of genes in the set (NGENES), effect size (positive values indicate enrichment) (BETA), standardized effect size (BETA_STD), standard error of the effect size (SE), p-value for enrichment (P), absolute value of the effect size beta (abs_beta), direction of the effect (beta_direction), adjusted p-value (BH) (adjP_val), -log10 of adjusted p-value ((minus) log10_adj_p).
